# The Role of Potassium and Calcium Currents in the Bistable Firing Transition

**DOI:** 10.1101/2023.08.16.553625

**Authors:** Fernando S. Borges, Paulo R. Protachevicz, Diogo L. M. Souza, Conrado F. Bittencourt, Enrique C. Gabrick, Lucas E. Bentivoglio, José D. Szezech, Antonio M. Batista, Iberê L. Caldas, Salvador Dura-Bernal, Rodrigo F. O. Pena

## Abstract

Healthy brains display a wide range of firing patterns, from synchronized oscillations during slow-wave sleep to desynchronized firing during movement. These physiological activities coexist with periods of pathological hyperactivity in the epileptic brain, where neurons can fire in synchronized bursts. Most cortical neurons are pyramidal regular spiking cells (RS) with frequency adaptation and do not exhibit bursts in current-clamp experiments (*in vitro*). In this work, we investigate the transition mechanism of spike-to-burst patterns due to slow potassium and calcium currents, considering a conductance-based model of a cortical RS cell. The joint influence of potassium and calcium ion channels on high synchronous patterns is investigated for different synaptic couplings (*g*_syn_) and external current inputs (*I*). Our results suggest that slow potassium currents play an important role in the emergence of high-synchronous activities, as well as in the spike-to-burst firing pattern transitions. This transition is related to bistable dynamics of the neuronal network, where physiological asynchronous states coexist with pathological burst synchronization. The hysteresis curve of the coefficient of variation of the inter-spike interval demonstrates that a burst can be initiated by firing states with neuronal synchronization. Furthermore, we notice that high-threshold (*I*_*L*_) and low-threshold (*I*_*T*_) ion channels play a role in increasing and decreasing the parameter conditions (*g*_syn_ and *I*) in which bistable dynamics occur, respectively. For high values of *I*_*L*_ conductance, a synchronous burst appears when neurons are weakly coupled and receive more external input. On the other hand, when the conductance *I*_*T*_ increases, higher coupling and lower *I* are necessary to produce burst synchronization. In light of our results, we suggest that channel subtype-specific pharmacological interactions can be useful to induce transitions from pathological high bursting states to healthy states.

## 1 Introduction

The classical Hodgkin-Huxley model, proposed in 1952, gave us a new perspective on understanding the electrical properties of neuronal dynamics considering the biophysical mechanics of the exchange of charges by sodium and potassium ion channels (1). Nowadays, much more is known about the variety of channels involved in the neuronal machinery (2; 3) and their relation to healthy brain function (4; 5) and diseases (6; 7). Despite that, the complete understanding of the role of such ion channels is still inciting many research questions in this matter (9; 8). Currently, an extensive perspective of research considering the neuroinformatics approach has provided new insights to understand and treat brain diseases (10; 11). Neuroinformatics has been used to fit and explore the dynamics of realistic neuron activities (12), including the influence of ion channels (13) and different currents to which neurons can be exposed (14; 15). In the present work, we are particularly interested in the ion channel influence as an adaptive mechanism that affects the firing pattern and synchronization transitions, since adaptation plays an important role in synchronization (16; 17).

The transition of neuronal single-cell firing patterns can emerge as a sum of different sources. In this framework, topology properties, such as the number (17; 18), type (19), the intensity of neuronal interactions (20; 21; 22; 23; 24), and the time delay of chemical transmission (25; 26) can contribute to the firing patterns transitions. Besides that, noise input currents, as well as external perturbations, can also play a role in the spike and burst transitions (27; 28; 29). The relationship of how single spikes and bursts are maintained in relation to theta and gamma oscillations was recently discovered (30). However, beyond the topology factors, intrinsic neuronal properties have been pointed out as the main factor in firing pattern transitions (32; 31). For example, the mechanism of adaptation of excitability appears to be a key factor in the transition between spike and burst patterns (33; 34). Typically, spike adaptation corresponds to the capability of neurons to reduce their spike frequency due to recent sub and over-threshold neuronal activities (35; 36). It is also known that potassium and calcium currents develop a role in neuronal adaptation mechanisms (37; 38; 39).

The emergence of different network firing patterns is mainly associated with intrinsic properties of the neurons, for example, ion channel density (32; 40) and type (42; 41), as well as the synaptic input currents (43; 44), noise (46; 45), and other kinds of couplings and external inputs that the neurons of the network can be submitted (47). A particular type of populational neuronal firing pattern in network dynamics is the synchronized state. In this context, the influence of the ion channels in synchronization was demonstrated by Boaretto et al. It was shown that small changes in ionic conductance can affect the capacity of the networks to exhibit phase synchronization (48). Landenbauer et al. studied the impact of adaptation on synchronization by comparing coupled neuronal models, and highlighting the importance of the comprehension of such mechanism on the emergence of the network dynamics (49). Synchronized activities are essential for the correct functioning of the brain; however, highly synchronized activities with burst firing patterns are associated with epileptic seizures (50). Besides the high synchronization levels, epileptic activities can also be related to burst firing patterns (51; 52) and low synaptic inhibition (53). The experimental results of cultured neuronal networks, based on multi-electrode arrays, indicated that burst activities present stronger synchrony capabilities with a sufficient level of excitation (54). These currents are also known to develop frequency-dependent resonant mechanisms, an example being the hyperpolarization-activated *I*_*h*_ current that is related to theta resonance and the T-type voltage-dependent Ca^2+^ channel that acts as an amplifier (55). In addition, T-type channels are related to the increase of burst activities in the seizure generation (56) and have been identified to play a central role in epileptogenesis in the pilocarpine model of epilepsy (57). Besides that, after status epilepticus, the density of T-type Ca2+ channels was upregulated in pyramidal neurons (58). However, a deeper understanding of the complementary roles of channels in the complex firing patterns of neurons and networks is still uncovered.

Bistable patterns are dynamical behaviors that can be exhibited from the levels of a single neuron (59) to a network (60; 61). Bistability in synchronization emerges as a collective dynamic interplay between excitatory and inhibitory interactions between neurons (62). Recently, Akcay et al. reported that bistability with different phases can emerge in a pair of Type I neurons connected by chemical synapses (63), a phenomenon related to the history dependency of the system and known as hysteresis. Bistability and hysteresis are two mechanisms that can be associated with the emergence of burst oscillations (64). In cortical neurons, the state-dependent in the coexistence of tonic and burst firings gather the conditions for the emergence of bistability and hysteresis (65). Simplified neuronal and oscillator models have reported the emergence of bistable firing patterns (33; 66). Bistable and multi-stable dynamics are said to play a role in both healthy (67) and abnormal brain activities (68). For this reason, studying a detailed description of ion channels in biophysical neuronal models can bring new light to understanding such firing patterns that emerge in brain activities.

The main purpose of this work is to investigate the impact of potassium and calcium currents as an adaptive mechanism that enables the emergence of burst synchronization associated with a bistable regime (33). In our simulations, burst activities emerge with highly synchronized firing patterns for a range of synaptic couplings (*g*_syn_) and external current input (*I*). Slow potassium currents promote the emergence of high-synchronous activities and spike-to-burst firing pattern transitions. This transition is bistable where physiological asynchronous states coexist with pathological burst synchronization. The hysteresis curve of the coefficient of variation of the inter-spike interval demonstrates that a burst can be initiated by synchronized firing states; however, asynchronous states result in an asynchronous spike firing pattern. Furthermore, we notice that for high values of high-threshold (*I*_*L*_) conductance, a synchronous burst appears when neurons are weakly coupled and receive more external input. On the other hand, when the conductance *I*_*T*_ increases, the opposite is observed. As our main conclusions, we show how the dynamic and biophysical characteristics of the slow potassium and calcium currents in networks promote bistable transitions. We believe that this work has the potential to uncover pharmacological targets in a manner in which high synchronization can be efficiently hindered.

Our paper is organized as follows. In Section 2, we describe the fundamental equations that govern the dynamics of the model and the diagnostics considered for the analysis of neuronal dynamics. In Section 3, we present our results. We depict neuronal dynamics by first presenting the perspective of a single network and, following, presenting to network configuration. We gradually introduce the role of potassium and calcium currents by observing how the emergency of the bistable regime takes place. Finally, in Section 4, we present our discussion of the expose our future perspectives and conclusions of this work.

## 2. Materials and Methods

### 2.1 Neural Model

We consider a conductance-based model in which the membrane potential *V* (70) is given by the following equation

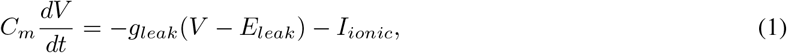

where *C*_m_ is the specific capacitance of the membrane, *g*_leak_ is the resting membrane conductance, *E*_leak_ the reversal potential, and *I*_*ionic*_ is the sum of partial ionic currents *I*_*j*_. The voltage-dependent ionic currents have the same general equation,

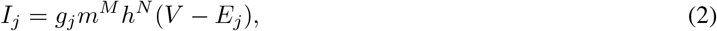

where the *j −* th ionic current *I*_*j*_ is expressed as the product of the maximum conductance of each ion *j* with the conductance *g*_*j*_ of the respective ion. The variables *m* and *n* are the activation variables of sodium and potassium, respectively, and *h* is the inactivation variable of the sodium ion channel (1). The difference between the membrane potential *V* and the reversal potential for a specific ion *E*_*j*_ is (*V − E*_*j*_) (70).

### 2.2 Description of ionic currents

The total ionic current *I*_*ionic*_ described in Equation 1 is given by

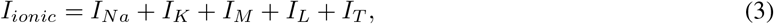

where *I*_*Na*_ and *I*_*K*_ are the basic sodium and potassium currents of the Hodgkin-Huxley model (71), *I*_*M*_ is a slow voltage-dependent potassium current responsible for spike-frequency adaptation (72), *I*_*L*_ is a high-threshold calcium current, and *I*_*T*_ is a low-threshold calcium current (73; 74).

#### 2.2.1 Sodium and Potassium currents

The voltage-dependent sodium and potassium currents are described by the Hodgkin-Huxley equations and were adapted for central neurons by Traub and Miles (71). The sodium currents are described in the following equations

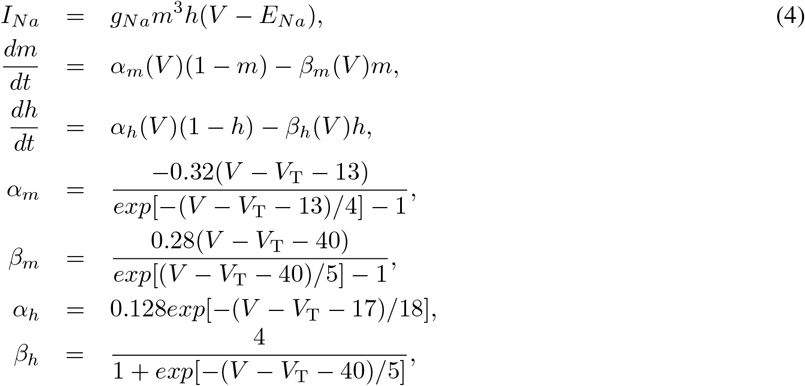

and the potassium currents are described by

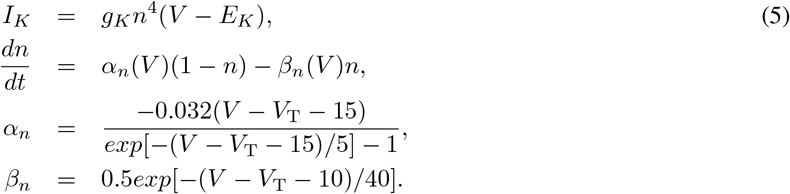

The conductance of sodium and potassium are *g*_*Na*_ = 50 mS/cm^2^, *g*_*K*_ = 5 mS/cm^2^, and the reversal potential are *E*_*Na*_ = 50 mV, *E*_*K*_ = *−* 100 mV, respectively. The *V*_T_ variable is used to adjust the peak threshold, in our simulations *V*_T_ = *−* 55 mV.

#### 2.2.2 Slow potassium current

The non-inactivating slow current of potassium ions is described by the equations

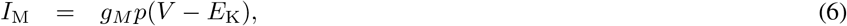

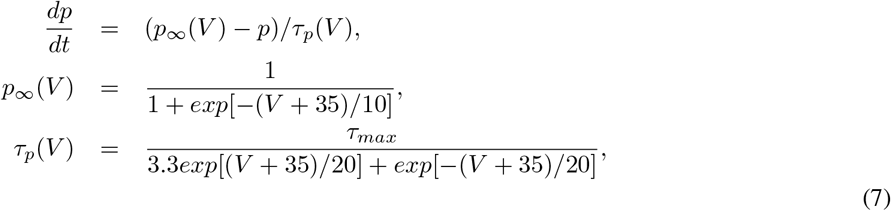

where *g*_*M*_ = 0.03 mS/cm^2^ and *τ*_max_ = 1000 ms (38; 72).

#### 2.2.3 Calcium currents

The first calcium current used to produce bursting is due to a high-threshold *Ca*^2+^ current described as

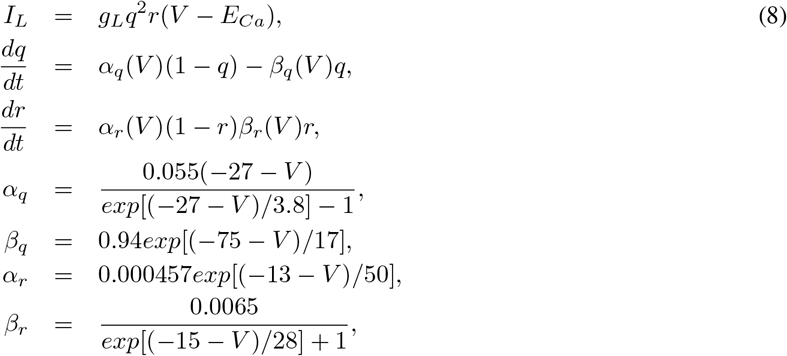

where the maximum conductance of *I*_L_ is *g*_*L*_ = 0.3 mS/cm^2^ (74).

The equations for the second type of calcium current, the low-threshold current, are

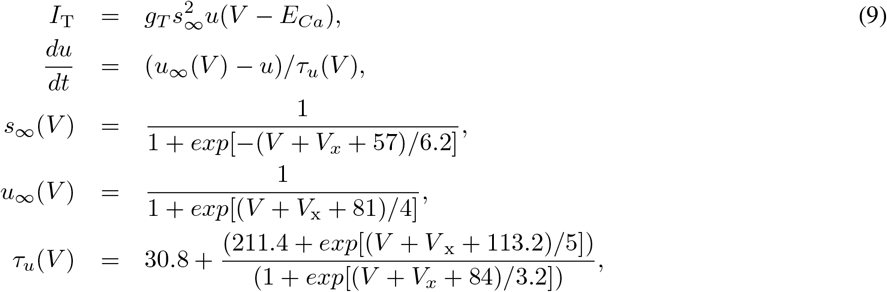

where *g*_*T*_ = 0.4 mS/cm^2^ is the maximal conductance of the low-threshold calcium current and *V* _x_ = 2 mV is a uniform shift of the voltage dependence (75; 76).

Calcium currents change intracellular calcium concentration [*Ca*^2+^]_*i*_, and as a consequence, the potential reversal of calcium ions (*E*_*Ca*_), which is given by

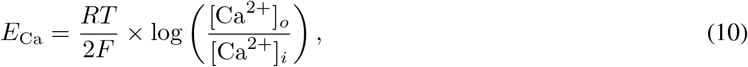

where *R* = 8.31 J *K*^*−* 1^*mol*^*−* 1^ is the universal gas constant, *T* = 309.15*K* is the temperature, *F* = 96485*Cmol*^*−* 1^ is the Faraday constant. The dynamics of the calcium concentration is given by

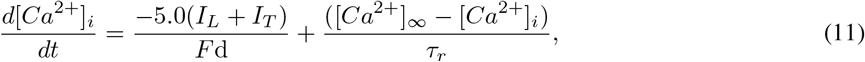

where [*Ca*^2+^]_*i*_ is intracellular *Ca*^2+^ concentration, [Ca^2+^]_*o*_ = 2.0 mM is extracellular *Ca*^2+^ concentration, [*Ca*^2+^]*∞* is the maximum concentration of calcium inside the cell, *d* = 1 *µ*m and *τ*_*r*_ = 5 ms.

#### 2.2.4 Neuronal network

We consider a randomly connected neuronal network composed of Hodgkin-Huxley neurons described in the previous sections, where the neurons are 80% excitatory and 20% inhibitory (77). We consider an Erdös–Rényi network with a connection probability *p* = 0.1, and there are no auto-connections (78). For the representation of each neuron *i* in a network, Eq. 1 with the addition of synaptic connections is represented by

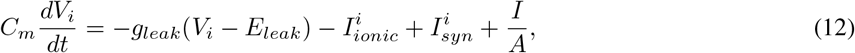

where *V* ^*i*^ represents the membrane potential, 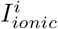 corresponds to the ion currents, *I* is a constant current equal to all neurons, and the superficial neuron area is *A* = *πdL*.

The chemical synaptic current that arrives in the neuron *i* is represented by

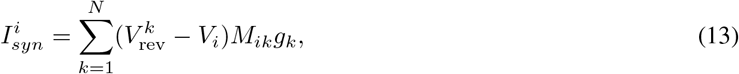

where 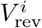 represents the synaptic reversal potential, *M* is the adjacency matrix of the connections, and *g*_*k*_ is the synaptic conductance from the neuron *k*. 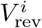 assumes value equal to 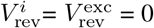 for excitatory and 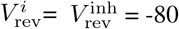 for inhibitory synaptic connection from the neuron *k*. The adjacency matrix is composed of element *M*_*ij*_ equal to 1 to represent connections from neuron *k* to neuron *i*, and equal to 0 to represent the absence of such connection. *g*_*k*_ is updated in the time of neuron *k* overpass *V* = 0 with a positive potential derivative (*dV/dt >* 0). The time of spikes of a certain neuron *k, t*_*k*_, is also defined by these two conditions in the membrane potential. In this way, the update in synaptic conductance is represented by *g*_*k*_*→ g*_*k*_ + *g*_syn_, where *g*_syn_ is the chemical intensity of synaptic updates, the same for excitatory and inhibitory connections. In addition to the update rule due to spike, each synaptic conductance *g*_*k*_ evolves by an exponential decay described by *dg*_*k*_*/dt* = *− g*_*k*_*/τ*_syn_ with *τ*_syn_ = 5.0 ms.

### 2.3 Measures

#### 2.3.1 Firing rate

We calculate the mean firing rate (in Hz) of all neurons in the network by

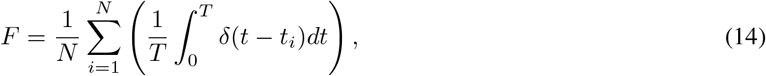

where *t*_*i*_ is the time of the *i*-th spike, N=1000 is the number of neurons, and *T* = 5 s is the time window considered for analyses.

#### 2.3.2 Coefficient of variation

We use the inter-spike intervals (ISI) where the *i*th interval is defined as the difference between two consecutive spike times *t*_*i*+1_ and *t*_*i*_, namely ISI_i_ = *t*_*i*+1_ *− t*_*i*_ *>* 0. From the ISI series, the first interval is referred to as ISI_1_ followed by the subsequent intervals, namely ISI_2_, ISI_3_, …, and ISI_n_. The ratio between the standard deviation and the mean (indicated by ⟨· ⟩) gives rise to the coefficient of variation

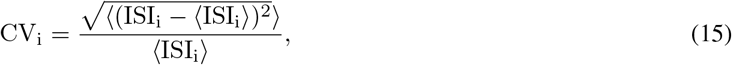

for the *ith* neuron. Finally, the average of CV_*i*_ over all neurons is given by

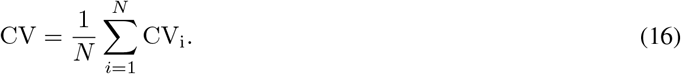

#### 2.3.3 Synchronization

To measure the level of synchronization exhibited by the neuronal network, we consider the complex order parameter of Kuramoto (79) given by

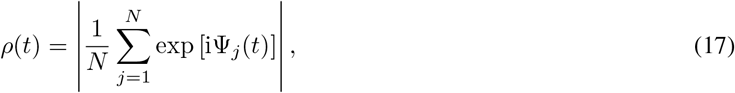

where the phase of each neuron *j* is represented by

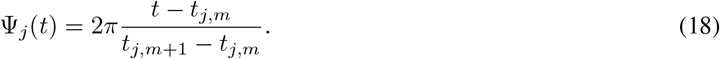

*t*_*j,m*_ represents the *m*-*th* spikes of the neuron *j*. The time *t* of the parameter is defined in the interval *t*_*j,m*_ *< t < t*_*j,m*+1_.

The time-average order parameter is calculated by

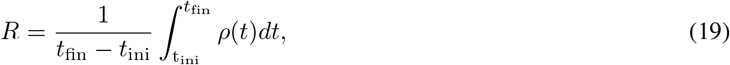

where *t*_ini_ and *t*_fin_ represent the initial and final time for the analyses of synchronization in the neuronal network (80).

The implementation of the numerical simulations was performed using self-developed C and NetPyNE codes (69) and can be freely accessed in Github.

## 3. Results

### 3.1 Neuron dynamics

We begin by presenting some essential dynamical characteristics of the neurons and networks. Fig. 1 displays the biophysical properties of neurons and their dependence on the ionic conductance of the slow potassium M-current and Fig. 2 with the additional presence of calcium currents.

**Figure 1:**
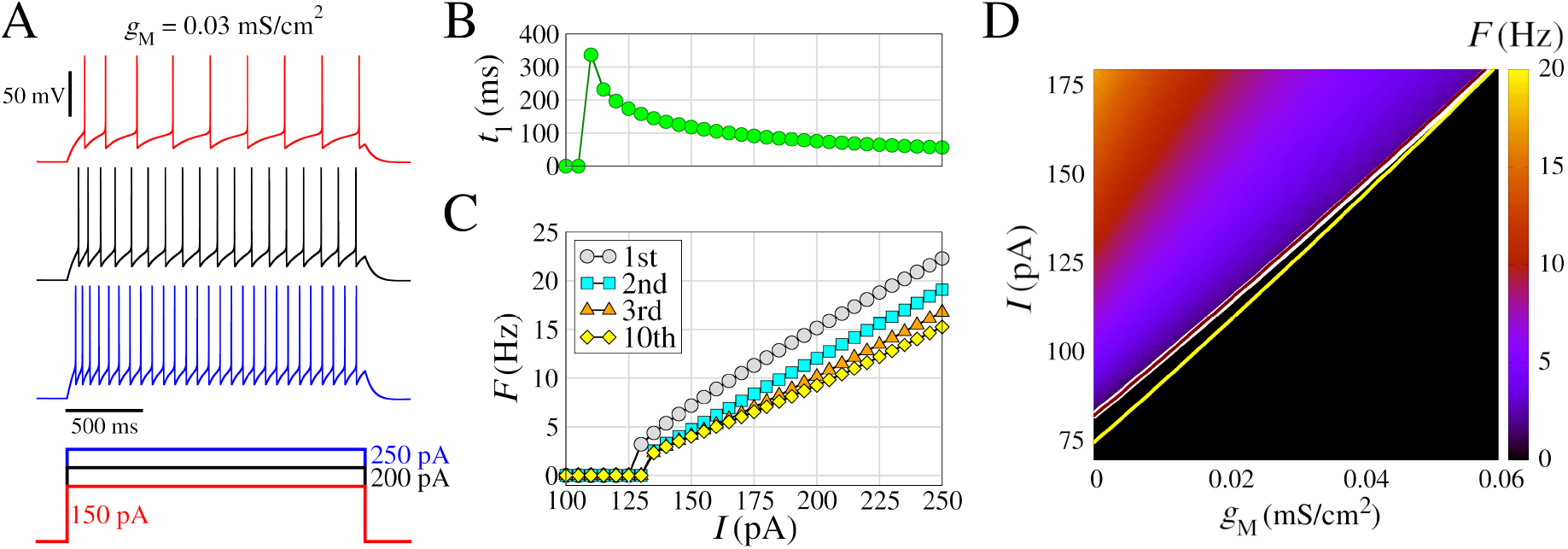
Model of regular spiking neuron, with *I*_*Na*_ **and** *I*_*K*_ for spike generation, and slow K+ current (*I*_*M*_) for spike-frequency adaptation. **(A)** (Top) Voltage traces with different amplitudes of the depolarizing pulses (bottom). **(B)** Time to first spike (*t*_1_) as a function of the injected current (amplitude of the pulse *I*). **(C)** Frequency-current curves (*F/I*), where the instantaneous firing rate (inverse of the inter-spike interval) is represented as a function of *I*. The curves indicated by different colors correspond to the 1st, the 2nd, the 3rd, and the 10th spike in the train. **(D)** Spike frequency *F* (in color) as function of *g*_*M*_ and *I*, considering 5.0 seconds time window. The white line represents the transition where *F >* 0 for *g*_T_ = 0 and *g*_L_ = 0. Additionally, this transition lines for *g*_T_ = 0 and *g*_L_ = 0.1 mS/cm^2^ (brown line), and *g*_T_ = 0.4 mS/cm^2^ and *g*_L_ = 0 (yellow line) are shown. Other parameters are *L* = *d* = 96.0 *µ*m, *g*_*leak*_ = 0.01 mS/cm^2^, *E*_*leak*_ = -85.0 mV, *g*_*Na*_ = 50 mS/cm^2^, *V*_*T*_ = *−* 55.0 mV, *g*_*K*_ = 5 mS/cm^2^, *τ*_max_ = 1000 ms, and *g*_*M*_ = 0.03 mS/cm^2^.

**Figure 2:**
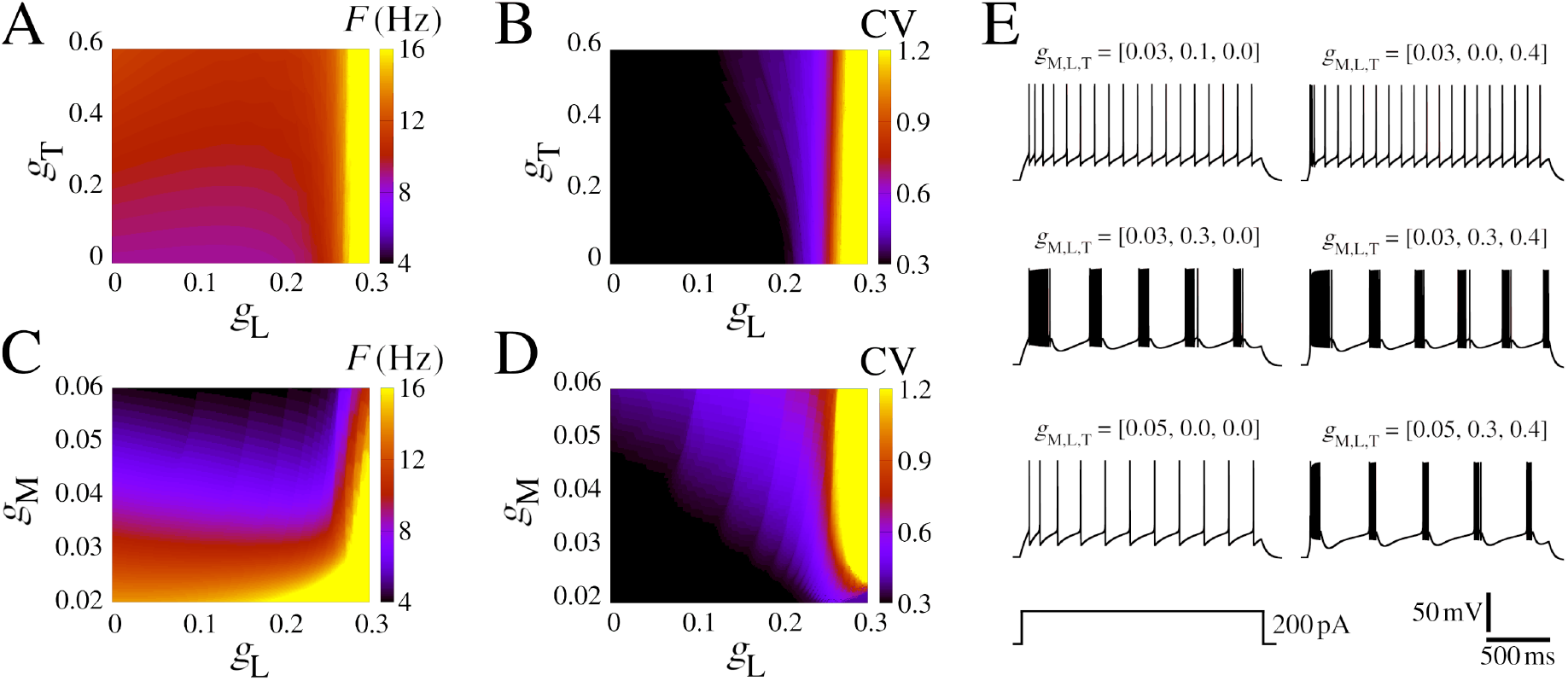
Firing pattern for different *I*_*M*_, *I*_*T*_, and *I*_*L*_ conductances. **(A)** Firing rate in colored (*g*_T_, *g*_L_)-diagram for *g*_M_ = 0.03 mS/cm^2^. **(B)** The same as **A** for the CV. **(C)** Firing rate in colored (*g*_M_, *g*_L_)-diagram for *g*_T_ = 0.4 mS/cm^2^. **(D)** The same as **C** for the CV. **(E)** Exemplar voltage traces considering different values of *g*_M_, *g*_L_, and *g*_T_, where each parameter combination is shown atop. Other parameters are the same as Fig. 1 with *I* = 200 pA.

In Fig. 1 we present extended characteristics for the regular spiking model by changing the input amplitudes. This model includes *I*_*Na*_ and *I*_*K*_ for the generation of spikes, and slow K^+^ current (*I*_*M*_) for the adaptation of the spike frequency. The frequency of the action potentials increases with the input amplitude, as shown by the three exemplar voltage traces (Fig. 1A). For input amplitudes of *I* = 110 pA, the neuron exhibits a single spike after*≈* 350 ms from the start of stimulus (Fig. 1B). The second spike occurs for *I* = 130 pA, where the first frequency *F* is obtained by 1*/*ISI_1_.

Regular spiking behavior is observed for *I >* 135 pA (Fig. 1C). As the amplitude increases, no bursting appears, but rather a linear increase in frequency with progressively lower frequencies for each spike in the train as a consequence of the adaptation mechanism (Fig. 1C).

In Fig. 1D we show *F* (in color) as a function of *g*_*M*_ and *I*, where *F* (Eq. 14) was calculated considering spikes in a time window of 5.0 seconds. The *g*_M_ has a great influence on the way the neuron fires by changing the minimum value of *I* where *F >* 0 (white line), in addition, higher values of *g*_M_ have lower values of *F* for the same value of *I*. This shows that the slow K^+^ current (*I*_*M*_), related to the adaptation of the spike frequency, has a strong influence on the response of the neuronal firing to external stimuli. On the other hand, low-threshold calcium currents (*I*_T_) have low alterations in *F* (yellow line) and high-threshold calcium currents (*I*_L_) have an almost null effect (brown line).

In order to understand the role of calcium currents in the neuron model, we show some diagrams that vary the low-threshold (*g*_T_) and high-threshold (*g*_L_) conductances. We delimited the range of the parameter based on the firing rate values lower than 20 Hz. Figs. 2A–D display colored (*g*_T_, *g*_L_) and (*g*_M_, *g*_L_)-diagrams for *F* and CV. Fig. 2E exhibits some examples of selected simulations. The initial burst extends into sustained bursting due to the influence of high-threshold Ca^2+^, this phenomenon becomes evident for *g*_L_ *>* 0.026 mS/cm^2^ (Fig. 2B). This observed frequency remains below 12 Hz for *g*_M_ *>* 0.03 mS/cm^2^ and *g*_L_ *<* 0.025 mS/cm^2^ (Fig. 2C), which is typically found in neurons in the rat somatosensory cortex (81). The coefficient of variation (CV) undergoes a transition from 0.3 to *>*1.0 depending on the value of *g*_L_ for *g*_M_ *>* 0.022 mS/cm^2^. This CV transition is abrupt and occurs at approximately *g*_L_ *≈*0.25 mS/cm^2^ (Fig. 2D). The two main effects of increases *g*_T_ are to reduce the time to the first spike and to generate the initial bursting pattern (Fig. 2E and Fig. S1). For some values of the *g*_L_ and *g*_M_ combination, regular bursts appear. Furthermore, the amplitude of the input current is also responsible for the change in the area in which regular burst activity (CV > 1.0) is observed in the colored diagram. Higher values of *I* increase the minimum *g*_M_ values to have bursts, for *I* = 200 pA the minimum *g*_M_ = 0.022 mS/cm^2^ while for *I* = 250 pA is necessary *g*_M_ *>* 0.038 mS/cm^2^ (Fig. S2).

The panels in Fig. 2 provide clear evidence of the impact of adaptation and burst development resulting from the influence of slow-potassium and calcium currents. In the next section, we will move forward with the network effects.

### 3.2 Neuron networks

Next, we study the behavior of networks of neurons, such as the ones discussed above, connected via chemical synapses. As such, the type of behavior varies depending on the input current *I* and the strength of the synapses *g*_syn_. A highly interesting phenomenon that arises in the network spiking pattern is the transition from asynchronous spiking to burst synchronization. Moreover, in this transition, bistable firing patterns are observed, where both asynchronous and synchronous states coexist.

In Fig. 3, we present a systematic study of the parameter combinations of constant applied current and the chemical synaptic conductance (*I* and *g*_syn_) in the measures *F*, CV, and *R* (see Sect. 2.3). These measures are displayed, respectively, in Fig. 3A, B, and C. Notice that here, we purposely remove the effect of *g*_M,L,T_ to show that without the slow potassium and calcium currents there is no burst no matter the input amplitude and coupling (Fig. 3B). Moreover, the firing rate increases with *I* and *g*_syn_ (Fig. 3A). Interestingly, however, as seen in Figs. 3C–D, even without the effect of *g*_M,L,T_ the raster plot shows a synchronized behavior with activity around 4 Hz due to population dynamics. This is noticed for combined low values of input current *I* and *g*_syn_ whereas for higher values only asynchronous behavior is observed (Fig. 3E).

**Figure 3:**
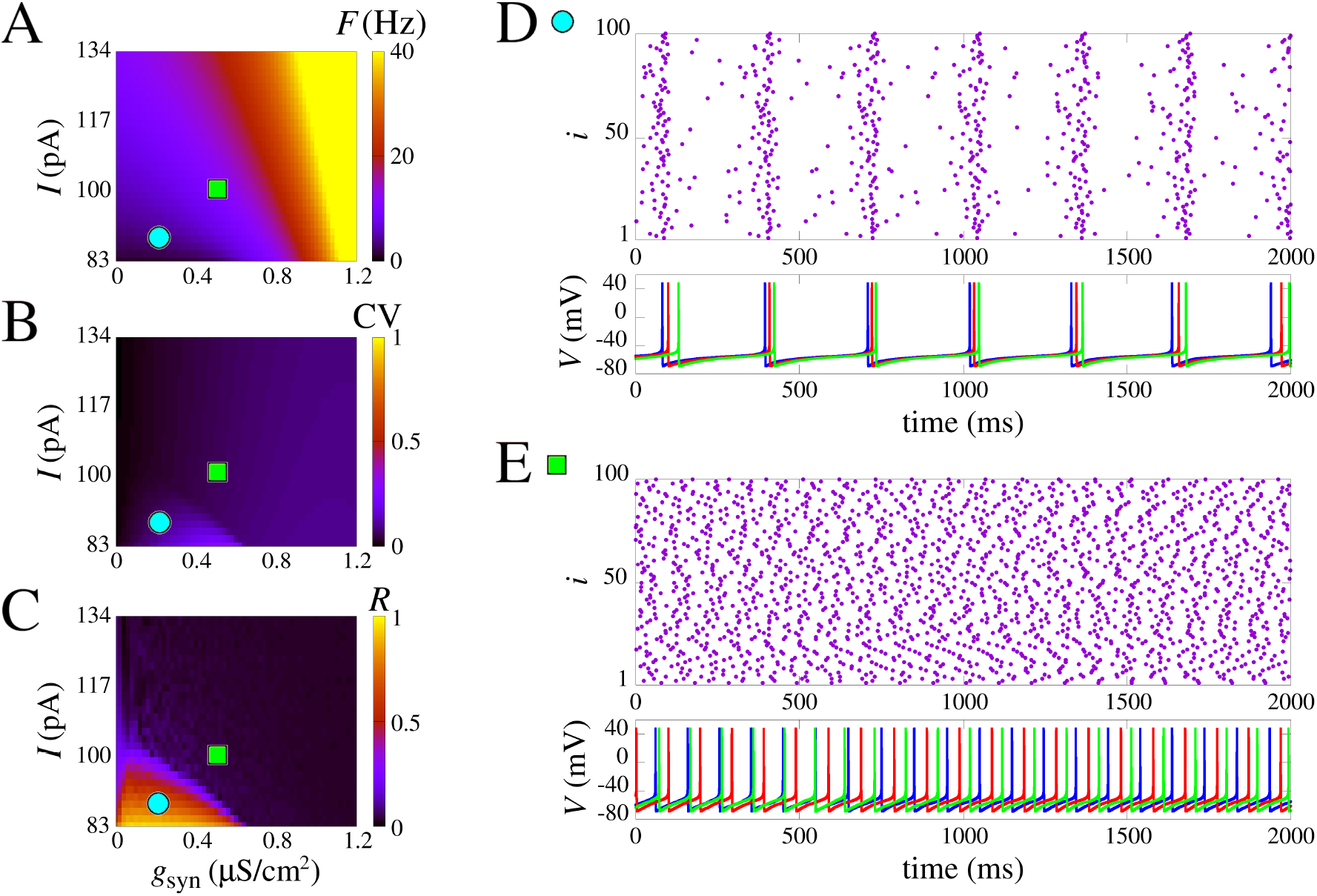
Spike synchronization in space parameters as a function of *I* and synaptic conductance (*g*_syn_). **(A)** Mean firing frequency. **(B)** Coefficient of variation, and **(C)** is the mean order parameter. **(D)** and **(E)** show the raster plot and the neuronal membrane potentials for the parameter indicated in **A**-**C. (D)** shows synchronized spikes for the parameters *I* = 88.3 pA and *g*_syn_= 0.2 *µ*S/cm^2^. **(E)** shows desynchronized spikes for the parameters *I* = 98.9 pA and *g*_syn_= 0.5 *µ*S/cm^2^. We use the model without slow potassium and calcium currents, i.e. *g*_M,L,T_ = [0, 0, 0].

The results demonstrate that the lack of slow potassium and calcium currents hinders the possibility of observing single-cell bursting even in a network. Transitions from asynchronous to synchronous activity are still present, as seen in how the circle and square are in areas of different values of *R* in Fig. 3C, but the CV changes are slight and nearly absent, indicating only spikes.

### Bistable regime

The results are different when the effect of the slow potassium current is added. As shown in Fig. 4, increasing *g*_syn_ results in transitions from asynchronous activity to a burst synchronization. Noticeably, the voltage traces exhibit individual bursting for neurons with a regular spike firing pattern without coupling. These network bursts are observed for fixed values of *I* < 200.0 pA and *g*_syn_ > 1.0 *µ*S/cm^2^.

**Figure 4:**
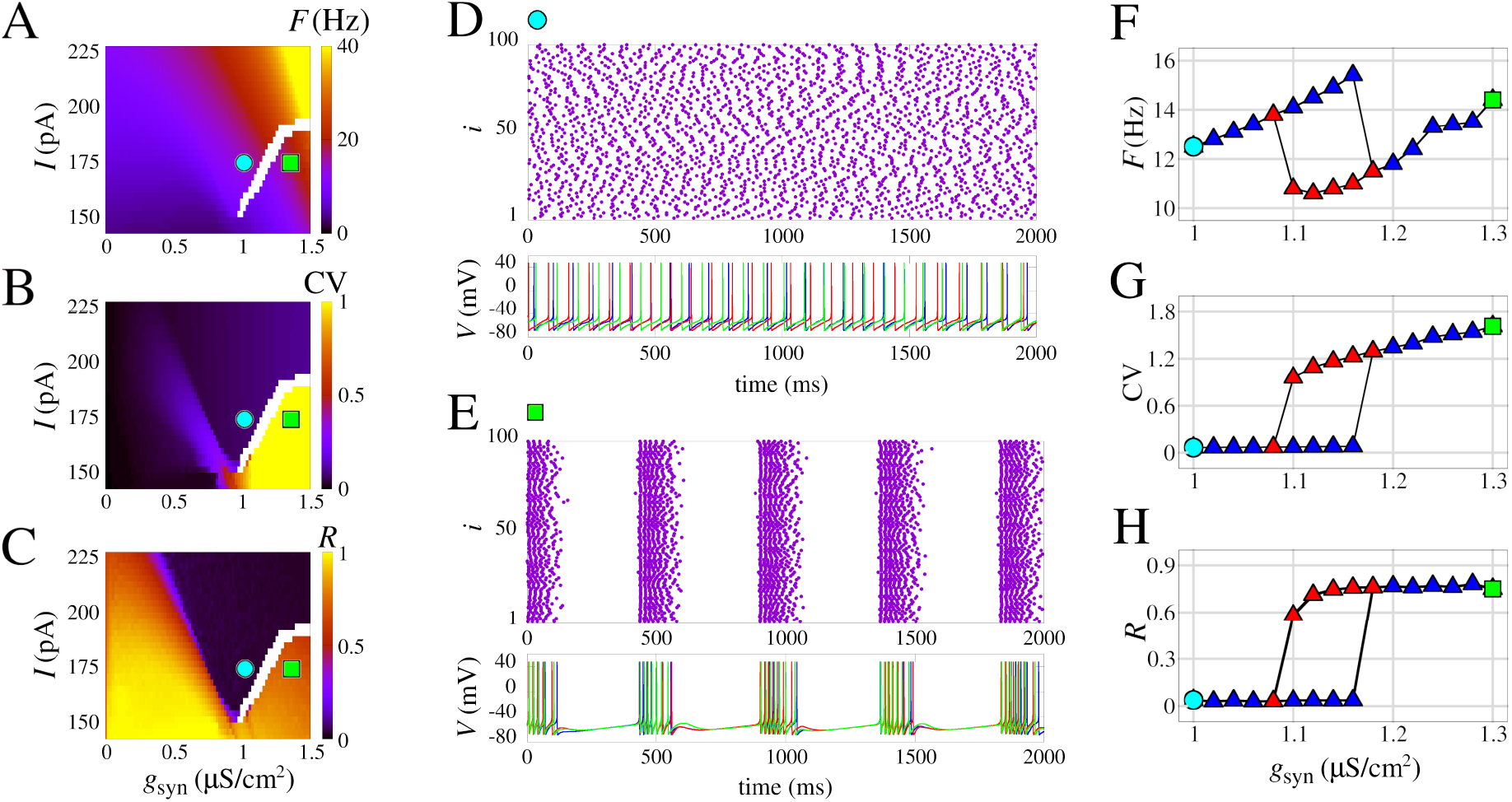
Burst synchronization in space parameters as a function of *I* and chemical synaptic conductance (*g*_syn_). **(A)** Mean firing frequency, **(B)** Coefficient of variation, and **(C)** Mean order parameter. **(D)** and **(E)** show the raster plot and the neuronal membrane potentials for the parameter indicated in **A**-**C**. In **(D)** desynchronized spikes for the parameters *I* = 170.4 pA and *g*_syn_= 1 *µ*S/cm^2^ are observed and synchronized bursts are shown for the parameters *I* = 170.4 pA and *g*_syn_= 1.3 *µ*S/cm^2^ in **E. (F–H)** Curves of *F*, CV, and *R*, respectively, for increasing values of *g*_syn_ (blue triangles) and decreasing values of *g*_syn_ (red triangles). We use the model with slow potassium and without calcium currents, i.e. *g*_M,L,T_ = [0.03, 0, 0] mS/cm^2^.

The emergence of the bursting synchronization by increasing values of *g*_syn_ has a marked effect on the measures *F*, CV, and *R*. When *I* = 170.4 pA, this sudden increase that leads to the transition starts approximately at *g*_syn_ = 1.09 *µ*S/cm^2^ for initial conditions with asynchronous spikes, and *g*_syn_ = 1.18 *µ*S/cm^2^ for initial conditions with burst synchronization. This hysteresis is indicated in the white area in Fig.4A–C and is an observation that is a strong indication of bistability. Fig. 4D shows a raster plot and voltage traces for the asynchronous spike pattern, and Fig. 4E shows the same for burst synchronization. A comparison of the blue and red curves in Fig. 4F–H shows the exact transition area from asynchronous activity to bursting synchronization.

The results demonstrate that the slow potassium current (here considered as *g*_M_ = 0.03 mS/cm^2^) can promote bistable dynamics in neuronal networks. This type of observation related to a biophysically grounded parameter is a key factor in understanding how bistability relates to other brain phenomena such as decision-making or pathologies such as epilepsy. The transition from spike to network burst was observed in (17) for the adaptative exponential integrate-and-fire neuron model, and, later, bistability in neuronal networks using this simple neuron model was related to epileptic seizures elsewhere (33; 90).

### Calcium effects in the bistability

We have shown that slow potassium current is necessary to observe the bistability. A natural question that arises from our work is how the firing transition and the bistable dynamics depend on the single-cell calcium ionic channels. In this section, we discuss how quantitative changes emerge for different combinations of calcium currents. Fig. 5 shows how CV values depend on constant current and chemical synaptic conductance (*I,g*_syn_) with respect to different values of *g*_M,L,T_. The bistable parameter region is identified in white and separate black and reddish/blueish regions for asynchronous spikes (low CV) to synchronous bursts (high CV), respectively. Fig. S3 demonstrates these qualitative patterns are maintained for the firing frequency *F* and the mean order parameter *R*.

**Figure 5:**
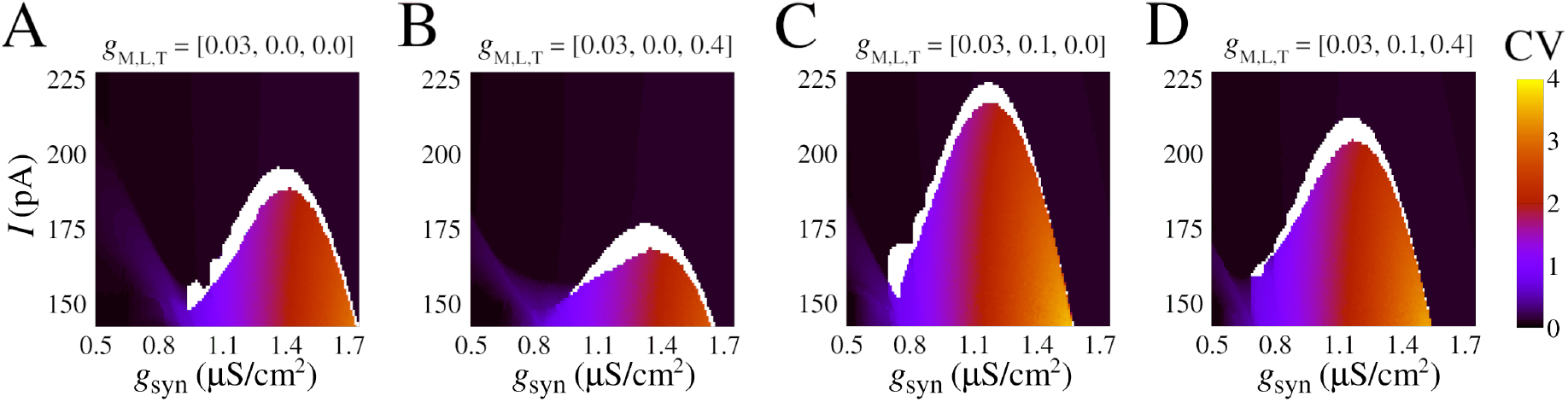
Effect of calcium currents in the burst synchronization in space parameters as a function of input current (*I*) and chemical synaptic conductance (*g*_syn_). **(A-D)** Coefficient of variation (CV) for different combinations of *g*_M,L,T_ (see values atop). **(A)** Model with slow potassium and without calcium currents, the same parameters of Fig. 4. **(B)** Model with slow potassium and only low-threshold calcium current. **(C)** Model with slow potassium and only high-threshold calcium current. **(D)** Model with slow potassium and both low-threshold and high-threshold calcium currents.

Indeed, the transition area changes according to the presence of ionic currents. The parameters *g*_syn_ and *I* are sensitive to changes with respect to *g*_M,L,T_. In particular, high-threshold calcium, controlled by *g*_L_, is the one that causes a greater change by allowing transitions at lower values of the *g*_syn_ (Fig. 5C). Furthermore, the synchronous area (reddish/blueish) is larger for the high-threshold calcium change, allowing for burst synchronization for *I >* 200.0 pA.

In contrast, low-threshold calcium promotes the opposite effect by slightly increasing the value of *g*_syn_ necessary to observe a transition (compare Fig. 5A to B and C to D).

Our results show that calcium has a double effect in promoting the bistability of asynchronous/synchronous activity that we observed: it can either facilitate it by allowing lower values of *g*_syn_ act on the transition when the high-threshold calcium channel is used or make it harder via an opposite effect mediated by the low-threshold calcium channel.

## 4 Discussion and Conclusion

In this work, we extend the analysis of bistable firing transitions in neurons and networks (61; 59; 60; 33) considering the effect of potassium and calcium currents. To quantify these firing patterns, we employed the usual measures, including firing rate, CV, and synchronization level. This provides an important link to the role of these ionic currents in controlling network behavior. We notice that in the absence of the joint effect of potassium and calcium, there is only a spike synchronization pattern and bursts at the single-cell level are not observed. In the presence of slow potassium and calcium currents, bistability is identified by hysteresis among synchronized bursting and asynchronous activity. It is worth noting a significant distinction from the study conducted by (33), which utilized the adaptative exponential integrate-and-fire neuron model. In our current model, we not only observe the effects but also establish a direct link to the underlying neurobiology, enabling us to propose experimental interventions that can be tested empirically through the manipulation of ionic currents.

Different voltage-dependent ion channels are responsible for the control of excitability and can play an intrinsic role in pathologies such as epilepsy (82; 83; 84), Parkinson (85), and Alzheimer’s disease (86). In particular, potassium and calcium ions have been found to play a role in many of these diseases (87; 88; 89). In addition, some research has evidenced that blocking such ion channels can play a role in avoiding the mechanism associated with the emergence of epileptic activities (91). Detailed investigations have been devoted to identifying the variations and mutations of calcium and potassium ion channels that exert the most influence on epileptic activity (92).

The opening and closing of such ion channels can have different effects on neuronal excitability. The majority of potassium ion channels open when the membrane depolarizes and close when it hyperpolarizes. The M current, in particular, acts in the subthreshold domain and limits the ability of the neuron to fire repetitively (93). Their involvement in epileptic activities is still being investigated (94; 95), and some recent research studying the lost- and gain-of-function of such ion has started to clarify these issues (96). Besides that, mutations in potassium channels have also been identified as an important factor in the parthenogenesis of human epilepsy (97). Such mutations and anomalies can be generated by external factors, e.g., the use of drug substances that have evidenced potential for changing the expression of potassium channels (98). Recently, precise studies and therapies have focused on the mutation of potassium channel genes (99). It is clearly important to further investigate how channel blocking mechanisms can serve to control network activity (100; 101).

Regarding calcium, we have shown that neurons embedded with these channels can exhibit a spike-to-burst transition (102). In particular, two types of calcium currents are highlighted: L-type and T-type. The former, where “L” stands for large or long-lasting, is a high-voltage activated channel. The latter, where “T” stands for transient, is a low-voltage activated channel. Calcium currents are highly involved in network high synchronous patterns that are observed in epileptic seizures (103). Not surprisingly, calcium channel blockers may act as anti-seizure drugs for prevention and treatment (105; 104), i.e., the blocking of such ion channels can attenuate burst firing pattern (106) and hinders epileptic depolarization (107). Nonetheless, it is challenging to develop drugs that are subtype-specific, and in most cases, they haven’t been developed yet (108). Thus, understanding the spike-to-burst pattern through computer simulations and neuroinformatics has the potential to elucidate the emergence of synchronous patterns related to epileptic activities (109). A combined treatment using calcium channels blockers was demonstrated to be beneficial in anticonvulsant and antinociceptive effects (110) (for the reader interested in channel blockers, please see the list in (108)). According to our results, whereas L-type channels make transitions easier by lowering the value of synaptic strength (*g*_syn_) necessary for the transition, T-type channels have the opposite effect.

Our results also extend the current knowledge of the joint effect of calcium and potassium ion channels in the context of other important firing pattern transitions. Based on our analysis, we can predict the ionic blockers required to avoid and treat high synchronous activity. These blockers would change the single-cell bursts that may result in epileptic seizures to asynchronous activity. In particular, we have evidence of the role of such ion channels not only in the spike-to-burst firing pattern transition but also in the relation of such transition with the synchronous patterns. Furthermore, we can gain insights by examining situations where the channels are blocked selectively, rather than all simultaneously.

## Funding acknowledge

The authors acknowledge the financial support from São Paulo Research Foundation (FAPESP, Brazil) (Grants N. 2018/03211-6, 2020/04624-2, 2021/12232-0, 2022/13761-9), Fundação Araucária and Coordenação de Aperfeiçoamento de Pessoal de Nível Superior - Brasil (CAPES), and NIH U24EB028998.

## Data availability

Numerical simulations and analysis were implemented using custom C and Python code, and using the NetPyNE modeling tool and the NEURON simulation engine, and can be freely accessed in Github. Data can be made available upon request.

## Supplementary Material

### S1 Neuron dynamics

Figure S1 illustrates the biophysical properties of neurons and their dependence on the ionic conductance of the slow potassium M-current with the additional presence of calcium currents. The frequency of action potentials increases with the input amplitude, as shown by the three exemplar voltage traces (Fig. S1A). For input amplitudes of *I* = 40 pA, the neuron presents a single spike after *≈*150 ms from the start of stimulus (Fig. S1B). The second spike occurs for *I* = 47 pA, where the first frequency *F* is obtained by 1*/*ISI_1_. Regular spiking behavior is observed for *I >* 160 pA (Fig. S1C). In Fig. S1D we show the minimum value of *I* where *F >* 0 for *g*_T_ = *g*_L_ = 0 (gray circles), the black line is a curve fit

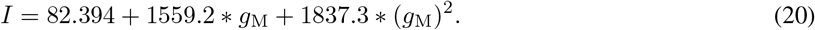

For *g*_T_ = 0.4 mS/cm^2^ and *g*_L_ = 0.2 mS/cm^2^ (orange squares) the fitted curve is

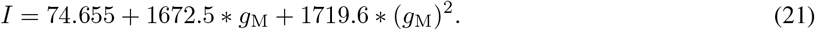

**Figure S1:**
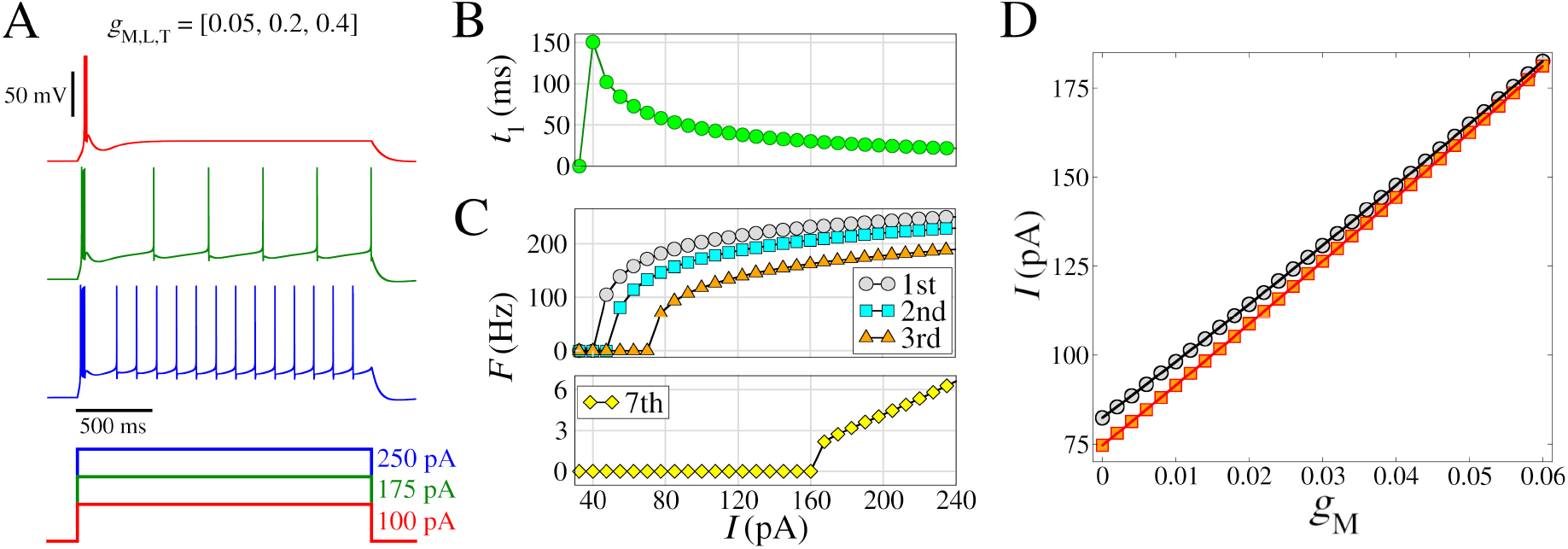
Model of regular spiking neuron, with *I*_*M*_, *I*_*T*_, and *I*_*L*_. **(A)** (Top) Voltage traces with different amplitudes of the depolarizing pulses (bottom). **(B)** Time to first spike (*t*_1_) as a function of the injected current (amplitude of the pulse *I*). **(C)** Frequency-current curves (*F/I*), where the instantaneous firing rate (inverse of the inter-spike interval) is represented as a function of *I*. The curves indicated by different colors correspond to the 1st, the 2nd, the 3rd, and the 7th spike in the train. **(D)** Transition where *F >* 0 for *g*_T_ = 0 and *g*_L_ = 0 (black line) and *g*_T_ = 0.4 mS/cm^2^ and *g*_L_ = 0.2 mS/cm^2^ (red line). Other parameters are *L* = *d* = 96.0 *µ*m, *g*_*leak*_ = 0.01 mS/cm^2^, *E*_*leak*_ = -85.0 mV, *g*_*Na*_ = 50 mS/cm^2^, *V*_*T*_ = *−* 55.0 mV, *g*_*Kd*_ = 5 mS/cm^2^, *τ*_max_ = 1000 ms, and *g*_*M*_ = 0.05 mS/cm^2^.

Figure S2 presents colored (*g*_M_, *g*_L_)-diagrams for *F* and CV when *I* = 200 pA (A–B) and *I* = 300 pA (C–D). The region in which sustained bursts occur (CV > 1.0) is lower for *I* = 300 pA than for *I* = 200 pA. The firing rate (*F*) increases when *I* = 300 pA for all (*g*_M_, *g*_L_)-diagram.

**Figure S2:**
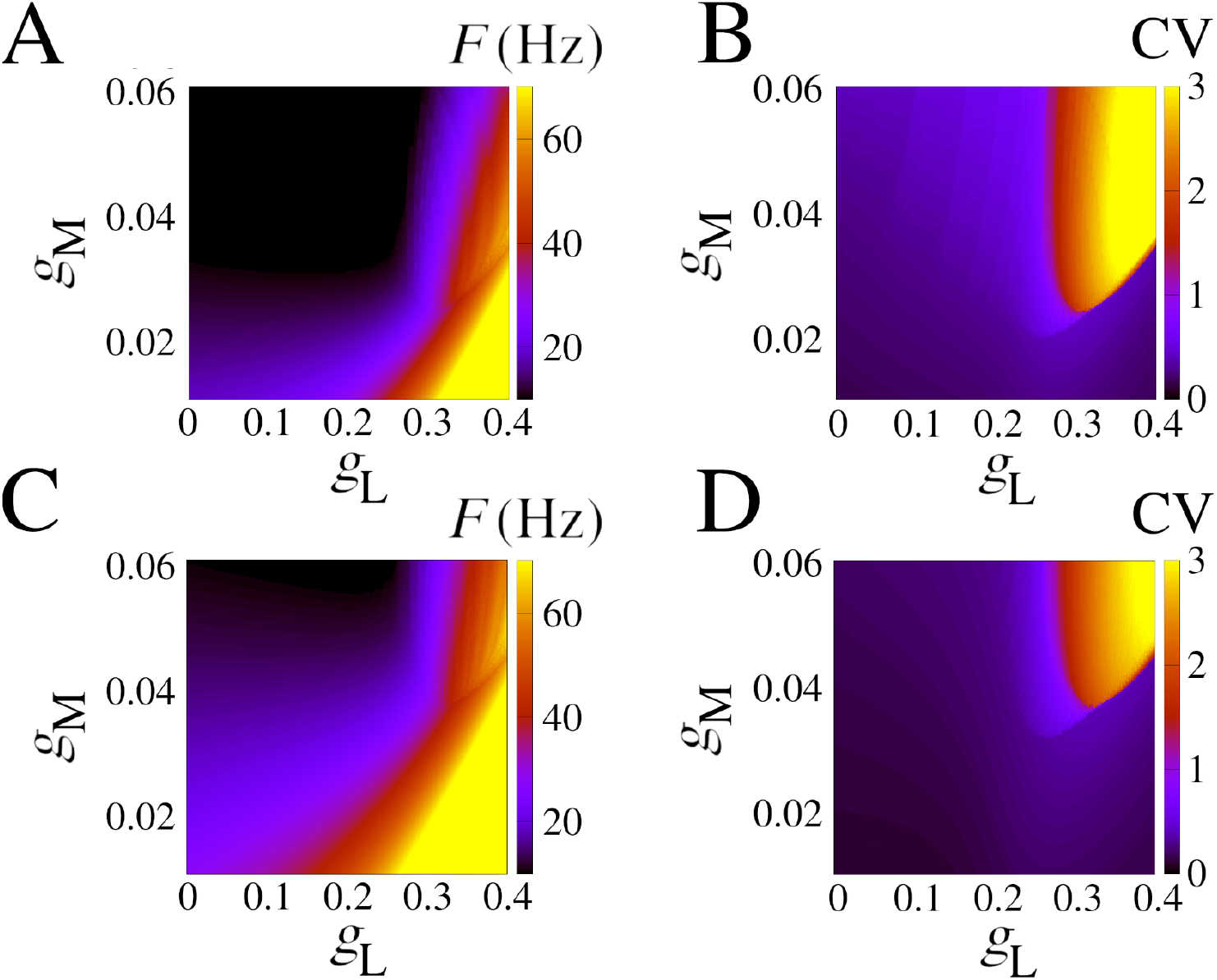
Firing pattern for diferent *I*_*M*_ and *I*_*L*_ conductances. **(A)** Firing rate in colored (*g*_M_, *g*_L_)-diagram for *I* = 200 pA. **(B)** The same as **A** for the CV. **(C)** Firing rate in colored (*g*_M_, *g*_L_)-diagram for *I* = 300 pA. **(D)** The same as **C** for the CV. Other parameters are the same as Fig. S1 with *g*_T_ = 0.4 mS/cm^2^.

### S2 Neuron network

In Figure S3 the values of CV, *F*, and *R* are shown at the top, middle, and bottom, respectively, in dependence on the constant current and the chemical synaptic conductance (*I,g*_syn_) with respect to different values of *g*_M,L,T_. The bistable parameter region is identified in white and separates the burst-synchronized region from the asynchronous ones. The high-threshold calcium conductance (*g*_L_) allows for transition at lower values of *g*_syn_ (Fig. S3C). Moreover, the burst-synchronized area is bigger in this case. In contrast, low-threshold calcium promotes the opposite effect by slightly increasing the value of *g*_syn_ necessary to observe a transition (compare Fig. S3A to B and C to D).

**Figure S3:**
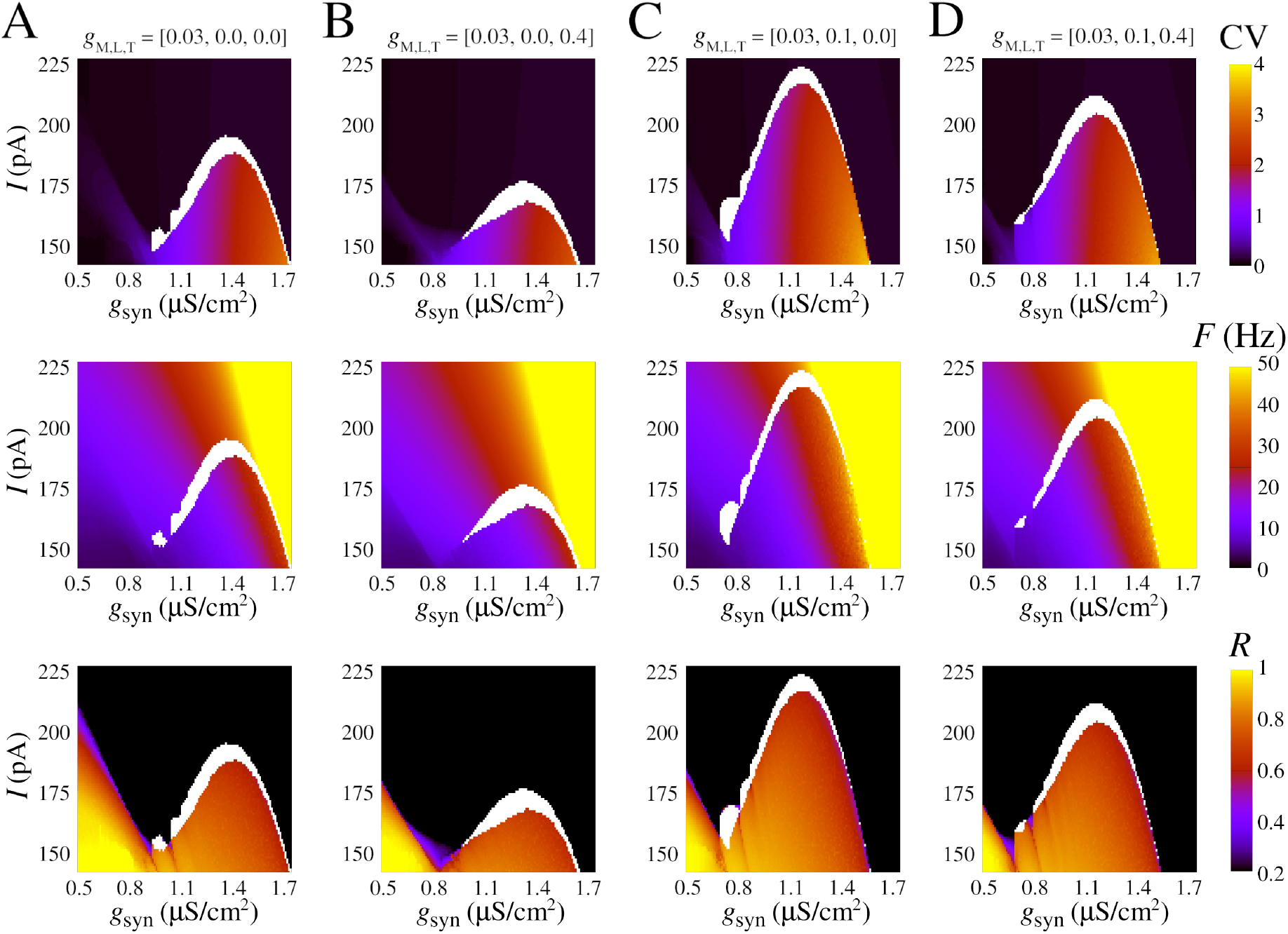
Changes in network firing pattern with input current (*I*) and chemical synaptic conductance (*g*_syn_). **(A-D)** CV (top), *F* (middle), and *R* (bottom) for different combinations of *g*_M,L,T_ (see values atop).

